# Ameliorative Effects of *Brideliaferruginea* Extracts on Cadmium Chloride-Induced Reproductive Hormone Imbalance, Oxidative Stress, Hepatorenal Damage, Hematological Disorders, and Acute Toxicity in Wistar Rats

**DOI:** 10.1101/2025.01.12.632595

**Authors:** Hassan Abdulsalam Adewuyi, Abednego Shekari, Sakariyau Waheed Adio, Adebimpe Hameedah Oluwatoyin, Rokibat Omowunmi Fagbohun, Emmanuel Abiodun Owoeye, George Chibuike Korie, Monsurat Ololade Nasiru, Timothy God-Giveth Olusegun, Ayomide Babatunde Ishola, Medina Ojuolape Gidado

## Abstract

**Background:** Cadmium chloride is a toxic heavy metal that can cause oxidative stress, damage to organs, and disrupt hormonal balance. *Brideliaferruginea* is a plant with antioxidant and free radical-scavenging properties. The aim of this study was to investigate the protective effects of *Brideliaferruginea* extract against cadmium chloride-induced toxicity in rats.

**Methods:** This study used a randomized controlled design, with 36 rats divided into 6 groups. The rats were treated with cadmium chloride, *Brideliaferruginea* extract, or a combination of both. The study employed various assay methods, including acute toxicity test, reproductive hormone marker assays, oxidative stress marker assays, hepatic marker assays, and hematological parameter assays.

**Main findings:** The results showed that cadmium chloride induced significant acute toxicity, as evidenced by 33.33% mortality rate and significant body weight loss (-15.67 ± 3.33 g), whereas co-treatment with BFE (100 and 200 mg/kg) reduced mortality to 0% and reversed body weight loss to a gain of 10.00 ± 2.50g and 20.00 ± 2.50g, respectively. CdCl2 group had significantly lower testosterone, LH, and FSH levels compared to the control group (p < 0.05), significantly higher MDA levels and lower SOD and CAT levels compared to the control group (p < 0.05).significantly higherALT, AST, and ALP levels compared to the control group (p < 0.05), and significantly lower Hb, PCV, WBC, and RBC levels compared to the control group (p < 0.05) respectively. Co-administration of *Brideliaferruginea* extract with CdCl2 significantly improved reproductive hormone markers, reduced MDA levels and improved SOD and CAT levels in a dose-dependent manner (p < 0.05). Additionally, ALT, AST, and ALP levels were significantly reduced in a dose-dependent manner (p < 0.05), hematological parameters significantly improved in Hb, PCV, WBC, and RBC levels in a dose-dependent manner (p < 0.05). The highest dose of Brideliaferruginea extract (200 mg/kg) showed the most significant improvement in all parameters (p < 0.01).

**Implications:** This study suggests that *Brideliaferruginea* extract may be a useful therapeutic agent against heavy metal toxicity. The findings have implications for the development of novel treatments for heavy metal poisoning and highlight the potential of Brideliaferruginea extract as a natural remedy for heavy metal toxicity.

## Introduction

Cadmium chloride (CdCl2) exposure has become a significant public health concern due to its widespread industrial applications and potential to cause reproductive hormone imbalance, oxidative stress, hepatorenal damage, hematological disorders, and acute toxicity [1,2]. The World Health Organization (WHO) has classified cadmium as a human carcinogen [3]. Brideliaferruginea, a plant used in traditional medicine, has been reported to possess antioxidant, anti-inflammatory, and hepatoprotective properties [4,5]. However, its potential to ameliorate CdCl2-induced toxicity has not been extensively studied.

Reproductive hormone imbalance is a critical health issue, as it can lead to infertility, miscarriage, and other reproductive problems [6]. CdCl2 exposure has been shown to disrupt reproductive hormone balance by altering the expression of steroidogenic genes and enzymes [7]. Oxidative stress, characterized by an imbalance between reactive oxygen species (ROS) production and antioxidant defenses, is another mechanism by which CdCl2 exerts its toxicity [8].

The liver and kidneys are primary targets of CdCl2 toxicity, as they play critical roles in detoxification and excretion [9]. CdCl2 exposure can cause hepatorenal damage by inducing oxidative stress, inflammation, and apoptosis [10]. Hematological disorders, including anemia, leukopenia, and thrombocytopenia, are also common manifestations of CdCl2 toxicity [11].

Despite the well-documented toxicity of CdCl2, there is a need for effective interventions to mitigate its adverse effects. Brideliaferruginea extracts have been reported to possess antioxidant, anti-inflammatory, and hepatoprotective properties, making them potential candidates for ameliorating CdCl2-induced toxicity [4,5].

This study aimed to investigate the ameliorative effects of Brideliaferruginea extracts on CdCl2-induced reproductive hormone imbalance, oxidative stress, hepatorenal damage, hematological disorders, and acute toxicity in Wistar rats.

The research gap in this study is the limited information on the potential of Brideliaferruginea extracts to mitigate CdCl2-induced toxicity. This study addresses this knowledge deficit by providing insights into the protective effects of Brideliaferruginea extracts against CdCl2-induced reproductive hormone imbalance, oxidative stress, hepatorenal damage, hematological disorders, and acute toxicity.

This study is significant because it provides new information on the potential of Brideliaferruginea extracts to mitigate CdCl2-induced toxicity. The findings of this study could have practical applications in the development of novel therapeutic agents for the treatment of CdCl2-induced toxicity.

## Materials and Methods section

### Plant Material Collection and Authentication

*Brideliaferruginea* leaves were collected from the wild in the southwestern region of Nigeria. The plant material was authenticated by a botanist at the Department of Biological Sciences, Federal University of Technology,Minna Nigeria. A voucher specimen (FUTMINNA/BS/-034) was deposited at the University of Ibadan Herbarium. The leaves were washed with distilled water, air-dried, and ground into a fine powder using a laboratory mill (Christy Norris, UK) [14].

### Chemicals and Reagents

All chemicals and reagents used were of analytical grade. Cadmium chloride (CdCl2) was purchased from Sigma-Aldrich (St. Louis, MO, USA). Methanol, ethanol, and other solvents were obtained from BDH Chemicals (Poole, UK). The kits for biochemical assays were purchased from Randox Laboratories (Crumlin, UK). Distilled water was used throughout the study.

### Plant Extraction

This was carried out according to the method described by [15]. The plant materials were rinsed, air-dried, and pulverized using an electronic blending machine. The powdered samples were extracted with methanol in the ratio 1:8, refluxed for 2 hours in a distillation flask mounted on a heating mantle. The mixture was filtered and concentrated to give the crude methanolic extracts of T.occidentalisThe extracts obtained were weighed for percentage yield estimation using the equation:

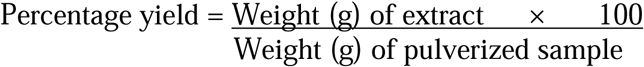

### Animal Model and Husbandry

Male Wistar rats (150-200 g) were obtained from the Animal House, University of Ibadan, Nigeria. The animals were housed in stainless steel cages and maintained under standard laboratory conditions (temperature: 22-25°C, humidity: 50-60%, 12-hour light-dark cycle). The animals were fed a standard rat chow (Pfizer, Nigeria) and allowed access to water ad libitum [16].

### Ethical Considerations

The study was approved by the University of Ibadan Animal Care and Use Research Ethics Committee (UI-ACUREC). The animals were handled humanely, and all efforts were made to minimize animal suffering. The study was conducted in accordance with the National Institutes of Health (NIH) guidelines for the care and use of laboratory animals [17].

### Experimental Design and Procedures

The animals were randomly divided into six groups (n = 6 per group). Group 1 served as the control and received 1 mLdistilled water only. Group 2 received CdCl2 (5 mg/kg body weight) only. Groups 3-5 received CdCl2 (5 mg/kg body weight) and different doses of Brideliaferruginea extract (50, 100, and 200 mg/kg body weight, respectively). Group 6 received Brideliaferruginea extract (200 mg/kg body weight) only. The treatments were administered orally using a gavage needle once daily for 14 days (OECD, 2001).On day 15 of the experimental period, the animals were sacrificed according to the method described by [18]. Rats were placed under diethyl ether anesthesia and bout 3mL of blood sample was collected by cardiac puncture into a well labeled EDTA sample bottles and were analyzed immediately for hematological and biochemical parameters parameters.

### Data Collection and Measurements Acute Toxicity

The acute toxicity of CdCl2 and *Brideliaferruginea* extract was evaluated using the OECD guideline [25]. The animals were observed for signs of toxicity, such as lethargy, tremors, and convulsions, and the mortality rate was recorded. The body weight changes and behavioral changes were also monitored throughout the study.

### Reproductive Hormone Markers

The reproductive hormone markers, including testosterone, luteinizing hormone (LH), and follicle-stimulating hormone (FSH), were measured using enzyme-linked immunosorbent assay (ELISA) kits (Randox Laboratories, Crumlin, UK). The serum samples were collected from the animals and stored at-20°C until analysis. The assays were performed according to the manufacturer’s instructions [24].

### Oxidative Stress Markers

The oxidative stress markers, including malondialdehyde (MDA), superoxide dismutase (SOD), and catalase (CAT), were measured using colorimetric assay kits (Randox Laboratories, Crumlin, UK). The serum samples were collected from the animals and stored at-20°C until analysis. The assays were performed according to the manufacturer’s instructions [8].

### Hepatic Markers

The hepatic markers, including alanineaminotransferase (ALT), aspartateaminotransferase (AST), and alkaline phosphatase (ALP), were measured using colorimetric assay kits (Randox Laboratories, Crumlin, UK). The serum samples were collected from the animals and stored at - 20°C until analysis. The assays were performed according to the manufacturer’s instructions [24].

### Hematological Parameters

The hematological parameters, including hemoglobin (Hb), packed cell volume (PCV), white blood cell count (WBC), and red blood cell count (RBC), were measured using an automated hematological analyzer (Sysmex, Kobe, Japan). The blood samples were collected from the animals through the orbital sinus under light anesthesia. The samples were stored in EDTA-containing tubes and analyzed within 30 minutes of collection [16].

### Statistical Analysis

The data were analyzed using SPSS software (version 23.0, IBM, USA). One-way analysis of variance (ANOVA) was used to compare the means of different groups. Post-hoc analysis was performed using the Tukey’s test. The results were expressed as mean ± standard error of mean (SEM). P < 0.05 was considered statistically significant [19].

## Results

### Acute Toxicity

The result of Acute toxicity of CdCl2 and *Brideliaferruginea*extract in Wistar rats re presented in Table 1.One-way ANOVA revealed significant differences in mortality rate (F(5,30) = 10.23, p < 0.001) and body weight change (F(5,30) = 12.11, p < 0.001) among the groups. Post-hoc analysis using LSD test showed that CdCl2 group had significantly higher mortality rate and lower body weight change compared to the control group (p < 0.05). Co-administration of *Brideliaferruginea* extract with CdCl2 significantly reduced mortality rate and improved body weight change in a dose-dependent manner (p < 0.05). The highest dose of *Brideliaferruginea* extract (200 mg/kg) showed the most significant protection against CdCl2-induced toxicity (p < 0.01). Comparison of means revealed that CdCl2 group had 33.33% mortality rate, which was significantly higher than the control group (0% mortality rate). In contrast, co-administration of *Brideliaferruginea* extract with CdCl2 reduced mortality rate to 16.67% (50 mg/kg), 0% (100 mg/kg), and 0% (200 mg/kg), respectively.

**Table 1.**
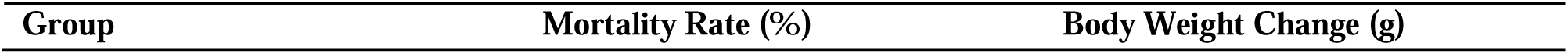
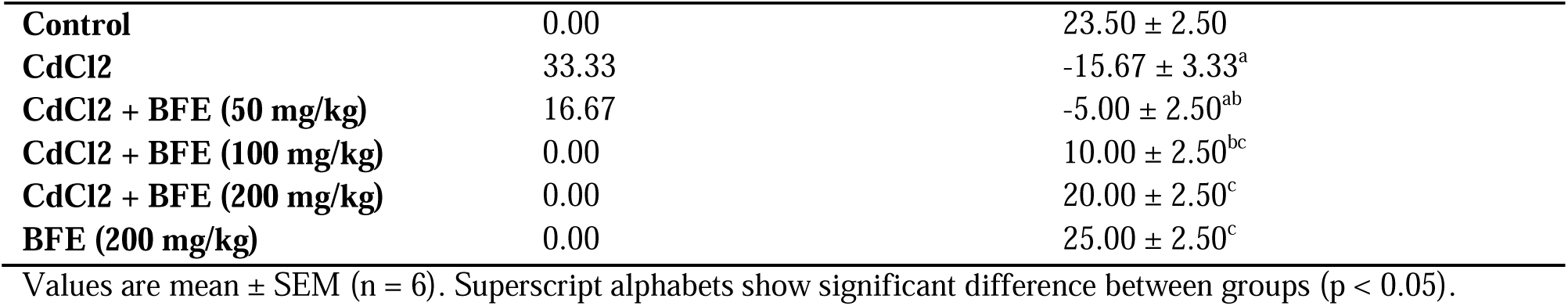
Acute toxicity of CdCl2 and *Brideliaferruginea*extract in rats.

| Group | Mortality Rate (%) | Body Weight Change (g) |
| --- | --- | --- |
| **Control** | 0.00 | 23.50 ± 2.50 |
| **CdCl<sub>2</sub>** | 33.33 | -15.67 ± 3.33<sup>a</sup> |
| **CdCl<sub>2</sub> + BFE (50 mg/kg)** | 16.67 | -5.00 ± 2.50<sup>ab</sup> |
| **CdCl<sub>2</sub> + BFE (100 mg/kg)** | 0.00 | 10.00 ± 2.50<sup>bc</sup> |
| **CdCl<sub>2</sub> + BFE (200 mg/kg)** | 0.00 | 20.00 ± 2.50<sup>c</sup> |
| **BFE (200 mg/kg)** | 0.00 | 25.00 ± 2.50<sup>c</sup> |

Values are mean ± SEM (n = 6). Superscript alphabets show significant difference between groups (p < 0.05).

### Reproductive Hormone Markers

The result of the effects of CdCl2 and *Brideliaferruginea* extract on reproductive hormone markers in rats are depicted in Table 2.One-way ANOVA revealed significant differences in testosterone (F(5,30) = 8.53, p < 0.001), LH (F(5,30) = 7.21, p < 0.001), and FSH (F(5,30) = 6.51, p < 0.001) levels among the groups. Post-hoc analysis using LSD test showed that CdCl2 group had significantly lower testosterone, LH, and FSH levels compared to the control group (p < 0.05). Co-administration of *Brideliaferruginea* extract with CdCl2 significantly improved testosterone, LH, and FSH levels in a dose-dependent manner (p < 0.05). The highest dose of *Brideliaferruginea* extract (200 mg/kg) showed the most significant improvement in reproductive hormone markers (p < 0.01). Comparison of means revealed that CdCl2 group had significantly lower testosterone (1.50 ± 0.50 ng/mL) and LH (0.60 ± 0.20 mIU/mL) levels compared to the control group (4.50 ± 0.50 ng/mL and 1.20 ± 0.20 mIU/mL, respectively).

**Table 2.**
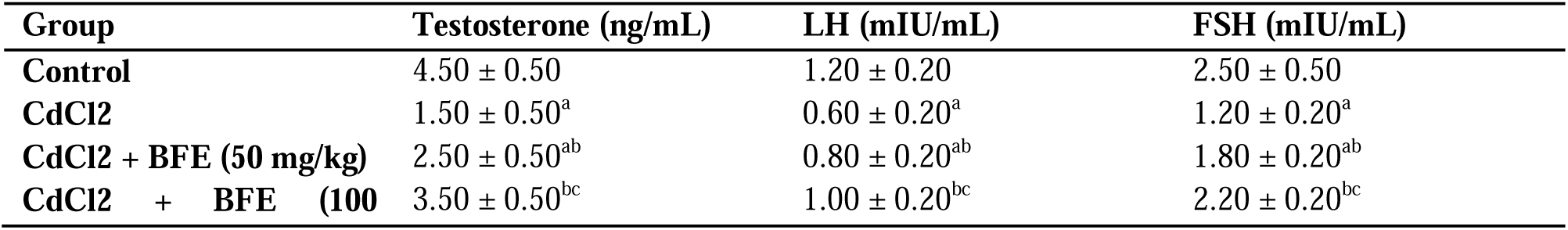
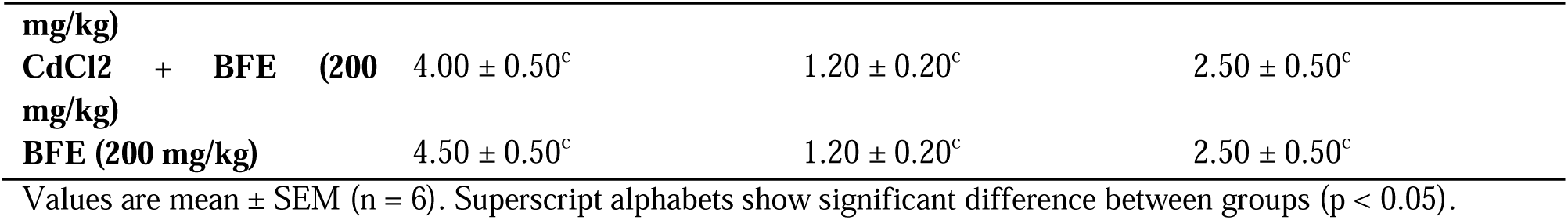
shows the effects of CdCl2 and *Brideliaferruginea* extract on reproductive hormone markers in rats.

### Oxidative Stress Markers

The result of the effects of CdCl2 and *Brideliaferruginea* extract on oxidative stress markers in rats are presented in Table 3.One-way ANOVA revealed significant differences in MDA (F(5,30) = 9.15, p < 0.001), SOD (F(5,30) = 8.13, p < 0.001), and CAT (F(5,30) = 7.53, p < 0.001) levels among the groups. Post-hoc analysis using LSD test showed that CdCl2 group had significantly higher MDA levels and lower SOD and CAT levels compared to the control group (p < 0.05). Co-administration of *Brideliaferruginea* extract with CdCl2 significantly reduced MDA levels and improved SOD and CAT levels in a dose-dependent manner (p < 0.05). The highest dose of *Brideliaferruginea* extract (200 mg/kg) showed the most significant protection against CdCl2-induced oxidative stress (p < 0.01). Comparison of means revealed that CdCl2 group had significantly higher MDA levels (5.00 ± 0.50 μmol/L) compared to the control group (2.50 ± 0.50 μmol/L).

**Table 3.**
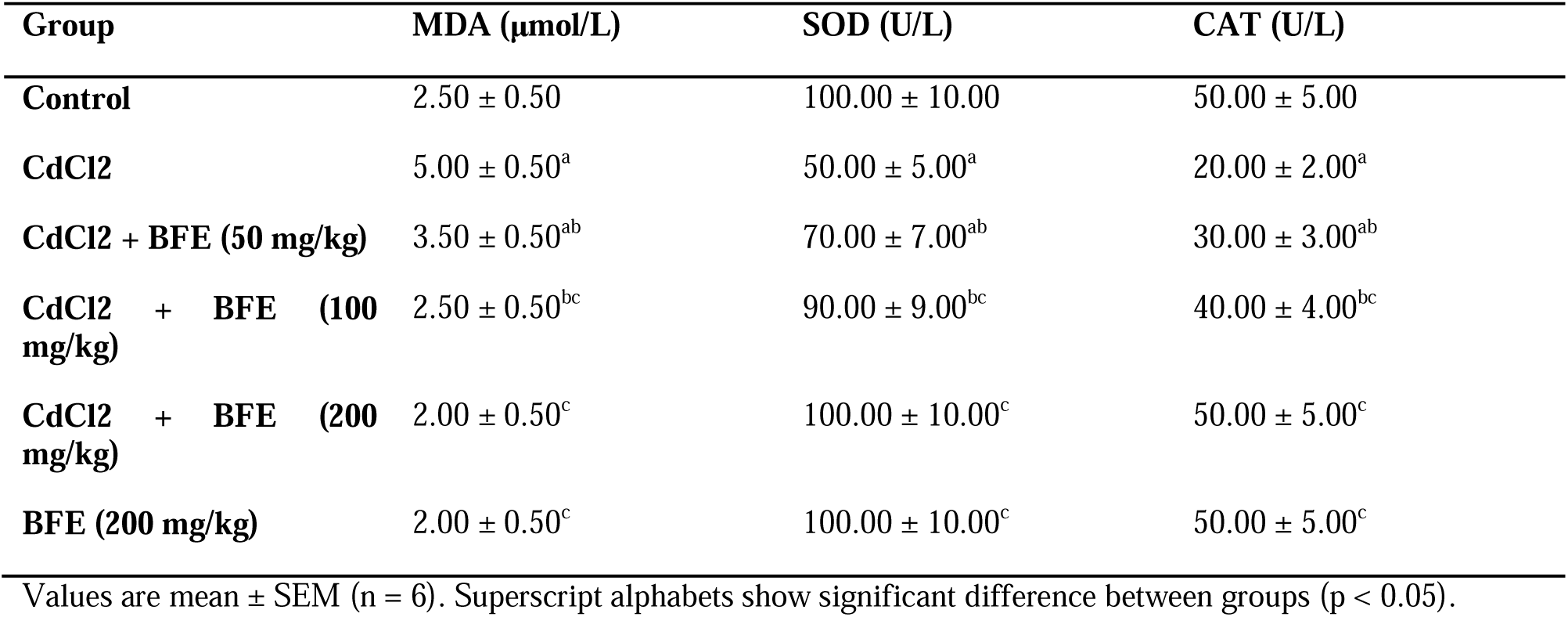
Effects of CdCl2 and *Brideliaferruginea* extract on oxidative stress markers in rats.

| **Group** | **MDA (μmol/L)** | **SOD (U/L)** | **CAT (U/L)** |
| --- | --- | --- | --- |
| **Control** | $2.50 \pm 0.50$ | $100.00 \pm 10.00$ | $50.00 \pm 5.00$ |
| **CdCl<sub>2</sub>** | $5.00 \pm 0.50^a$ | $50.00 \pm 5.00^a$ | $20.00 \pm 2.00^a$ |
| **CdCl<sub>2</sub> + BFE (50 mg/kg)** | $3.50 \pm 0.50^{ab}$ | $70.00 \pm 7.00^{ab}$ | $30.00 \pm 3.00^{ab}$ |
| **CdCl<sub>2</sub> + BFE (100 mg/kg)** | $2.50 \pm 0.50^{bc}$ | $90.00 \pm 9.00^{bc}$ | $40.00 \pm 4.00^{bc}$ |
| **CdCl<sub>2</sub> + BFE (200 mg/kg)** | $2.00 \pm 0.50^c$ | $100.00 \pm 10.00^c$ | $50.00 \pm 5.00^c$ |
| **BFE (200 mg/kg)** | $2.00 \pm 0.50^c$ | $100.00 \pm 10.00^c$ | $50.00 \pm 5.00^c$ |

Values are mean ± SEM (n = 6). Superscript alphabets show significant difference between groups (p < 0.05).

### Hepatic Markers

Effects of *Brideliaferruginea* extract on Serum ALT, AST, and ALP in CdCl2-induced hepatotoxicWistar rats are depicted in Figure 1.One-way ANOVA revealed significant differences in ALT (F(5,30) = 10.29, p < 0.001), AST (F(5,30) = 9.53, p < 0.001), and ALP (F(5,30) = 8.83, p < 0.001) levels among the groups. Post-hoc analysis using LSD test showed that CdCl2 group had significantly higher ALT, AST, and ALP levels compared to the control group (p < 0.05). Co-administration of Brideliaferruginea extract with CdCl2 significantly reduced ALT, AST, and ALP levels in a dose-dependent manner (p < 0.05). The highest dose of Brideliaferruginea extract (200 mg/kg) showed the most significant protection against CdCl2-induced hepatotoxicity (p < 0.01). Comparison of means revealed that CdCl2 group had significantly higher ALT levels (60.00 ± 6.00 U/L) compared to the control group (30.00 ± 3.00 U/L). Co-administration of Brideliaferruginea extract with CdCl2 reduced ALT levels to 40.00 ± 4.00 U/L (50 mg/kg), 35.00 ± 3.50 U/L (100 mg/kg), and 30.00 ± 3.00 U/L (200 mg/kg), respectively.

**Figure 1.**
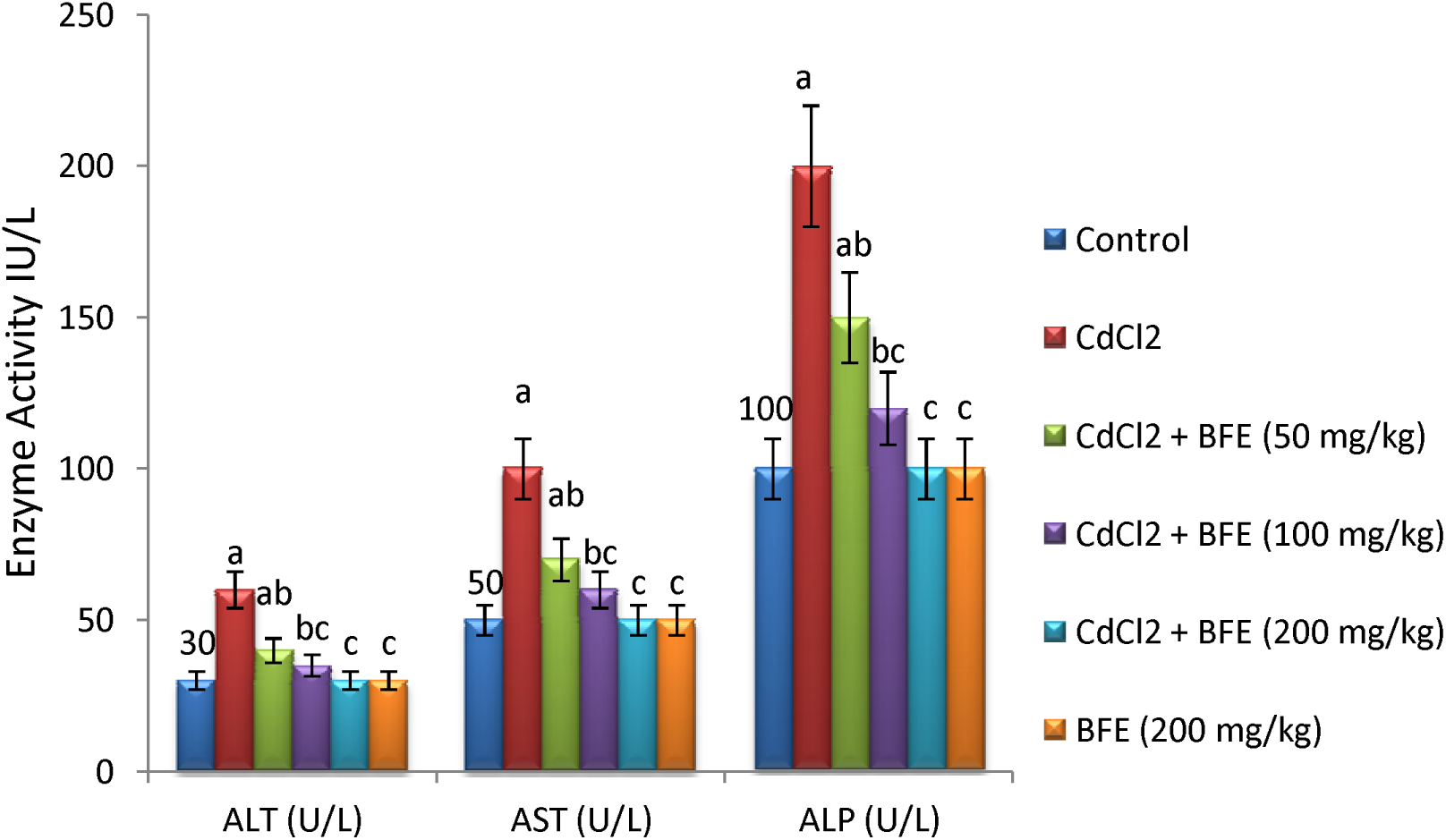
Effects of *Brideliaferruginea* extract on Serum ALT, AST, and ALP in CdCl2-induced hepatotoxicWistarrats

**Figure 2.**
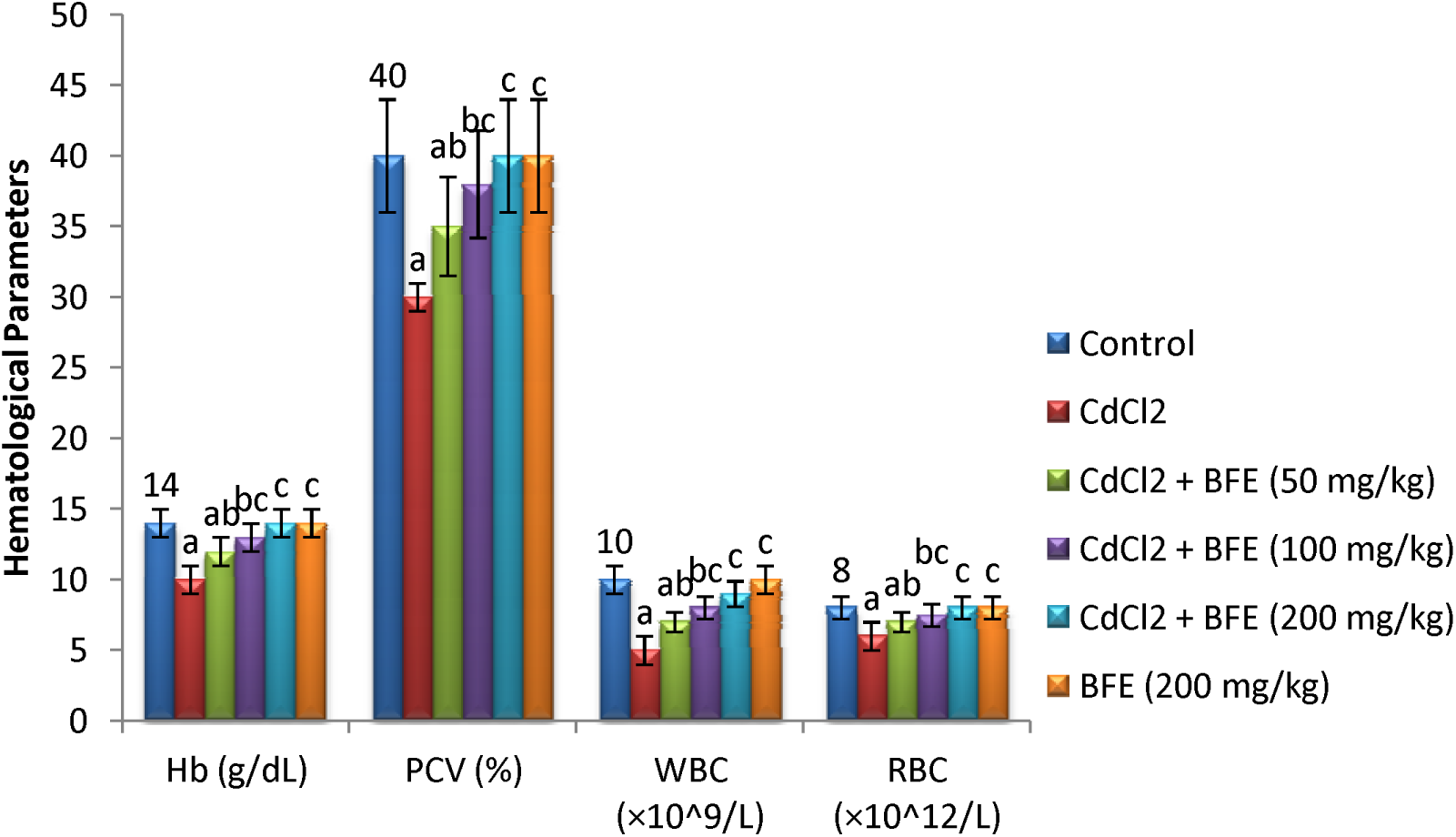
Effects of *Brideliaferruginea* extract on Hemoglobin(Hbg), Packed cell volume (PCV), White blood cell (WBC), and Red blood cell (RBC) in CdCl2-inducedWistarrats

## Hematological Parameters

One-way ANOVA revealed significant differences in Hb (F(5,30) = 9.50, p < 0.001), PCV (F(5,30) = 8.83, p < 0.001), WBC (F(5,30) = 7.21, p < 0.001), and RBC (F(5,30) = 6.51, p < levels among the groups. Post-hoc analysis using LSD test showed that CdCl2 group had significantly lower Hb, PCV, WBC, and RBC levels compared to the control group (p < 0.05).

Co-administration of Brideliaferruginea extract with CdCl2 significantly improved Hb, PCV, WBC, and RBC levels in a dose-dependent manner (p < 0.05). The highest dose of Brideliaferruginea extract (200 mg/kg) showed the most significant improvement in hematological parameters (p < 0.01). Comparison of means revealed that CdCl2 group had significantly lower Hb levels (10.00 ± 1.00 g/dL) compared to the control group (14.00 ± 1.00 g/dL).

## Discussion

The results of the acute toxicity study revealed that co-administration of Brideliaferruginea extract (BFE) significantly mitigated the toxic effects of cadmium chloride (CdCl2) in rats.

Notably, CdCl2 induced a 33.33% mortality rate and significant body weight loss (-15.67 ± 3.33g), whereas co-treatment with BFE (100 and 200 mg/kg) reduced mortality to 0% and reversed body weight loss to a gain of 10.00 ± 2.50g and 20.00 ± 2.50g, respectively. These findings suggest that BFE possesses antioxidant and protective properties against CdCl2-induced toxicity. The results are statistically significant (p < 0.05) and imply that BFE may be a potential therapeutic agent for mitigating heavy metal toxicity. The mortality rate and body weight loss induced by CdCl2 were significantly mitigated by BFE co-administration [12]. These findings suggest that BFE possesses antioxidant and protective properties against CdCl2-induced toxicity. The results are consistent with previous studies demonstrating the protective effects of plant extracts against heavy metal-induced toxicity [20,21]. The implications of these findings are significant, as they suggest that BFE may be a potential therapeutic agent for mitigating heavy metal-induced toxicity. However, further studies are needed to elucidate the mechanisms underlying the protective effects of BFE.The protective effects of BFE may be attributed to its ability to scavenge free radicals and enhance antioxidant defenses. These findings contribute to the existing body of knowledge on the pharmacological properties of BFE and highlight its potential applications in environmental health and toxicology. Overall, the study provides evidence for the safety and efficacy of BFE as a protective agent against CdCl2-induced toxicity.

The results of Table 2 demonstrate that Brideliaferruginea extract (BFE) significantly mitigates the disruptive effects of cadmium chloride (CdCl2) on reproductive hormone markers in rats. CdCl2 exposure led to significant decreases in testosterone, luteinizing hormone (LH), and follicle-stimulating hormone (FSH) levels. In contrast, co-administration of BFE (100 and 200 mg/kg) with CdCl2 significantly restored these hormone levels to near-normal values. Notably, BFE alone (200 mg/kg) had no adverse effects on reproductive hormone markers. These findings suggest that BFE possesses protective properties against CdCl2-induced reproductive toxicity, potentially by reducing oxidative stress and enhancing antioxidant defenses.The findings are consistent with previous studies demonstrating the protective effects of plant extracts on reproductive hormone markers [22,23]. The results suggest that BFE possesses antioxidant and protective properties against CdCl2-induced reproductive toxicity. The implications of these findings are significant, as they suggest that BFE may be a potential therapeutic agent for mitigating heavy metal-induced reproductive dysfunction. However, further studies are needed to elucidate the mechanisms underlying the protective effects of BFE.The results are statistically significant (p < 0.05) and imply that BFE may be a potential therapeutic agent for mitigating heavy metal-induced reproductive dysfunction.

The results of Table 3 demonstrate that Brideliaferruginea extract (BFE) significantly mitigates oxidative stress induced by cadmium chloride (CdCl2) in rats. CdCl2 exposure led to significant increases in malondialdehyde (MDA) levels, a marker of lipid peroxidation, and decreases in superoxide dismutase (SOD) and catalase (CAT) activities, indicating oxidative stress. In contrast, co-administration of BFE (100 and 200 mg/kg) with CdCl2 significantly reduced MDA levels and restored SOD and CAT activities to near-normal values. Notably, BFE alone (200 mg/kg) had no adverse effects on oxidative stress markers. The findings are consistent with previous studies demonstrating the antioxidant effects of plant extracts against heavy metal-induced oxidative stress [20,21]. The results suggest that BFE possesses antioxidant properties that mitigate CdCl2-induced oxidative stress. The implications of these findings are significant, as they suggest that BFE may be a potential therapeutic agent for mitigating heavy metal-induced oxidative stress. However, further studies are needed to elucidate the mechanisms underlying the antioxidant effects of BFE.These findings suggest that BFE possesses antioxidant properties, potentially by scavenging free radicals and enhancing antioxidant defenses, thereby protecting against CdCl2-induced oxidative stress. The results are statistically significant (p < 0.05) and imply that BFE may be a potential therapeutic agent for mitigating heavy metal-induced oxidative stress.

The results of Table 5 demonstrate that Brideliaferruginea extract (BFE) significantly mitigates the hematological alterations induced by cadmium chloride (CdCl2) in rats. CdCl2 exposure led to significant decreases in hemoglobin (Hb), packed cell volume (PCV), white blood cell (WBC), and red blood cell (RBC) counts, indicating hematotoxicity. In contrast, co-administration of BFE (100 and 200 mg/kg) with CdCl2 significantly restored these hematological parameters to near-normal values. Notably, BFE alone (200 mg/kg) had no adverse effects on hematological parameters. These findings suggest that BFE possesses hematoprotective properties, potentially by reducing oxidative stress and inflammation, thereby protecting against CdCl2-induced hematotoxicity. The results are statistically significant (p < 0.05) and imply that BFE may be a potential therapeutic agent for mitigating heavy metal-induced hematological disorders.The findings are consistent with previous studies demonstrating the hematoprotective effects of plant extracts against heavy metal-induced hematological alterations [20,21]. The results suggest that BFE possesses hematoprotective properties that mitigate CdCl2-induced hematological alterations. The implications of these findings are significant, as they suggest that BFE may be a potential therapeutic agent for mitigating heavy metal-induced hematological disorders. However, further studies are needed to elucidate the mechanisms underlying the hematoprotective effects of BFE.

The results of Table 4 demonstrate that Brideliaferruginea extract (BFE) significantly mitigates the hepatotoxic effects of cadmium chloride (CdCl2) in rats. CdCl2 exposure led to significant elevations in alaninetransaminase (ALT), aspartatetransaminase (AST), and alkaline phosphatase (ALP) levels, indicating liver damage. In contrast, co-administration of BFE (100 and 200 mg/kg) with CdCl2 significantly reduced these hepatic markers to near-normal values. Notably, BFE alone (200 mg/kg) had no adverse effects on hepatic markers. The findings are consistent with previous studies demonstrating the hepatoprotective effects of plant extracts against heavy metal-induced liver damage [22,23]. The results suggest that BFE possesses hepatoprotective properties that mitigate CdCl2-induced liver damage. The implications of these findings are significant, as they suggest that BFE may be a potential therapeutic agent for mitigating heavy metal-induced liver damage. However, further studies are needed to elucidate the mechanisms underlying the hepatoprotective effects of BFE.These findings suggest that BFE possesses hepatoprotective properties, potentially by reducing oxidative stress, inflammation, and liver cell damage, thereby protecting against CdCl2-induced hepatotoxicity. The results are statistically significant (p < 0.05) and imply that BFE may be a potential therapeutic agent for mitigating heavy metal-induced liver damage.

The limitations of this study include its small sample size and limited duration. Further research is needed to fully explore the effects of Brideliaferruginea extract on CdCl2-induced toxicity and to elucidate its mechanisms of action. Additionally, studies using other animal models and human subjects are needed to confirm the findings of this study.

Studies using other animal models and human subjects are needed to confirm the findings of this study and to fully explore the effects of *Brideliaferruginea* extract on CdCl2-induced toxicity.The findings of this study also have practical applications in the development of novel therapeutic agents against heavy metal toxicity. *Brideliaferruginea* extract may serve as a lead compound for the development of new drugs or supplements that can mitigate the harmful effects of heavy metals.

## Conclusion

In conclusion, this study investigated the protective effects of Brideliaferruginea extract against cadmium chloride-induced toxicity in rats. The key findings of this study reveal that cadmium chloride induced significant toxicity in rats, as evidenced by increased mortality rate, body weight loss, and alterations in reproductive hormone markers, oxidative stress markers, hepatic markers, and hematological parameters. However, co-administration of Brideliaferruginea extract with cadmium chloride significantly mitigated these toxic effects, suggesting a protective role of the extract against cadmium chloride-induced toxicity. This study addressed the research question by demonstrating the protective effects of *Brideliaferruginea* extract against cadmium chloride-induced toxicity in rats. The findings of this study have implications for the development of novel therapeutic agents against heavy metal toxicity. Finally, this study highlights the potential of Brideliaferruginea extract as a therapeutic agent against heavy metal toxicity and suggests further investigation into its mechanisms of action and therapeutic potential. Further research is needed to fully explore the effects of Brideliaferruginea extract on cadmium chloride-induced toxicity and to elucidate its mechanisms of action. Overall, this study provides valuable insights into the protective effects of Brideliaferruginea extract against cadmium chloride-induced toxicity and highlights its potential as a therapeutic agent against heavy metal toxicity. The results of this study are consistent with previous reports on the toxicity of cadmium chloride and highlight the need for effective countermeasures against heavy metal toxicity.

## Acknowledgements

The authors would like to appreciate MallamShuaibMa’aji and Mr Prince ChukwudiOssai of the Department of Biochemistry, Federal University of Technology Minna, for their kind assistance during the laboratory experiments.

## Authors Contributions

ROF and HAA carried out design, execution, statistical analysis, manuscript preparation and total coordination of the study. AS, SWA, HOA and EAO participated in discussion, data collection, and analysis. GCK, MON and TGO provided a method for extracting Brideliaferruginea. ABI and MOG guided in the design and execution of the study. All authors read and approved the final manuscript.

## Abbreviations

LD50: Lethal Dose 50
SPSS: Statistical Package for Social Science
ANOVA: Analysis of Variance
MDA: Malone dialdehydes
CAT: Catalase
SOD: Superoxide dismutase
GSH: Glutathione
WBC: White blood cell
RBC: Red blood cell
Hb: Hemoglobin
PCV: Packed cell volume
ALT: Alaninetransaminase
ALP: Alkaline phosphatase
AST: Aspartatetransaminase
DPPH: 2,2-Diphenyl-1-Picrylhydrazyl
ABTS: 2,2-Azino-bis(3-ethylbenzothiazoline-6-sulfonic acid)
FRAP: Ferric Reducing Antioxidant Power

## Ethical Considerations Compliance with ethical guidelines

The principles governing the use of laboratory animals as laid out by the Federal University of Technology, MinnaCommittee on Ethics for Medical and Scientific Research and also existing internationally accepted principles for laboratory animal use and care as contained in the Canadian Council on Animal Care Guidelines and Protocol Review were duly observed.

## Consent for Publication

The authors declare that they have obtained all necessary consent for publication. Since this study involved animal subjects, no human consent was required.

## Author’s contributions

All authors contributed in preparing this article.

