## Supplemental tables 1, 2 and 3 for "Ameliorative Effects of *Brideliaferruginea* Extracts on Cadmium Chloride-Induced Reproductive Hormone Imbalance, Oxidative Stress, Hepatorenal Damage, Hematological Disorders, and Acute Toxicity in Wistar Rats"

**Table 1. Acute toxicity of CdCl2 and *Brideliaferruginea*extract in rats**

| Group | Mortality Rate (%) | Body Weight Change (g) |
| --- | --- | --- |
| Control | 0.00 | 23.50 ± 2.50 |
| CdCl2 | 33.33 | -15.67 ± 3.33^a^ |
| CdCl2 + BFE (50 mg/kg) | 16.67 | -5.00 ± 2.50^ab^ |
| CdCl2 + BFE (100 mg/kg) | 0.00 | 10.00 ± 2.50^bc^ |
| CdCl2 + BFE (200 mg/kg) | 0.00 | 20.00 ± 2.50^c^ |
| BFE (200 mg/kg) | 0.00 | 25.00 ± 2.50^c^ |

Values are mean ± SEM (n = 6). Superscript alphabets show significant difference between groups (p < 0.05).

**Table 2 shows the effects of CdCl2 and *Brideliaferruginea* extract on reproductive hormone markers in rats**

| Group | Testosterone (ng/mL) | LH (mIU/mL) | FSH (mIU/mL) |
| --- | --- | --- | --- |
| Control | 4.50 ± 0.50 | 1.20 ± 0.20 | 2.50 ± 0.50 |
| CdCl2 | 1.50 ± 0.50^a^ | 0.60 ± 0.20^a^ | 1.20 ± 0.20^a^ |
| CdCl2 + BFE (50 mg/kg) | 2.50 ± 0.50^ab^ | 0.80 ± 0.20^ab^ | 1.80 ± 0.20^ab^ |
| CdCl2 + BFE (100 mg/kg) | 3.50 ± 0.50^bc^ | 1.00 ± 0.20^bc^ | 2.20 ± 0.20^bc^ |
| CdCl2 + BFE (200 mg/kg) | 4.00 ± 0.50^c^ | 1.20 ± 0.20^c^ | 2.50 ± 0.50^c^ |
| BFE (200 mg/kg) | 4.50 ± 0.50^c^ | 1.20 ± 0.20^c^ | 2.50 ± 0.50^c^ |

Values are mean ± SEM (n = 6). Superscript alphabets show significant difference between groups (p < 0.05).

**Table 3. Effects of CdCl2 and *Brideliaferruginea* extract on oxidative stress markers in rats.**

| Group | MDA (μmol/L) | SOD (U/L) | CAT (U/L) |
| --- | --- | --- | --- |
| Control | 2.50 ± 0.50 | 100.00 ± 10.00 | 50.00 ± 5.00 |
| CdCl2 | 5.00 ± 0.50^a^ | 50.00 ± 5.00^a^ | 20.00 ± 2.00^a^ |
| CdCl2 + BFE (50 mg/kg) | 3.50 ± 0.50^ab^ | 70.00 ± 7.00^ab^ | 30.00 ± 3.00^ab^ |
| CdCl2 + BFE (100 mg/kg) | 2.50 ± 0.50^bc^ | 90.00 ± 9.00^bc^ | 40.00 ± 4.00^bc^ |
| CdCl2 + BFE (200 mg/kg) | 2.00 ± 0.50^c^ | 100.00 ± 10.00^c^ | 50.00 ± 5.00^c^ |
| BFE (200 mg/kg) | 2.00 ± 0.50^c^ | 100.00 ± 10.00^c^ | 50.00 ± 5.00^c^ |

Values are mean ± SEM (n = 6). Superscript alphabets show significant difference between groups (p < 0.05).

Enzyme Activity IU/L

|  | Control | CdCl2 | CdCl2 + BFE (50 mg/kg) | CdCl2 + BFE (100 mg/kg) | CdCl2 + BFE (200 mg/kg) | BFE (200 mg/kg) |
| --- | --- | --- | --- | --- | --- | --- |
| ALT (U/L) | 30 | 60 | 40 | 35 | 30 | 30 |
| AST (U/L) | 50 | 100 | 70 | 60 | 50 | 50 |
| ALP (U/L) | 100 | 200 | 150 | 120 | 100 | 100 |

**Figure 1. Effects of *Brideliaferruginea* extract on Serum ALT, AST, and ALP in CdCl2-induced hepatotoxicWistarrats**

Hematological Parameters

|  | Control | CdCl2 | CdCl2 + BFE (50 mg/kg) | CdCl2 + BFE (100 mg/kg) | CdCl2 + BFE (200 mg/kg) | BFE (200 mg/kg) |
| --- | --- | --- | --- | --- | --- | --- |
| Hb (g/dL) | 14 | 10 | 12 | 13 | 14 | 14 |
| PCV (%) | 40 | 30 | 35 | 38 | 40 | 40 |
| WBC (×10^9/L) | 10 | 5 | 7 | 8 | 9 | 10 |
| RBC (×10^12/L) | 8 | 6 | 7 | 7.5 | 8 | 8 |

**Figure 2.Effects of *Brideliaferruginea* extract on Hemoglobin(Hbg), Packed cell volume (PCV), White blood cell (WBC), and Red blood cell (RBC) in CdCl2-inducedWistarrats**
